# Temporal assessment of fire severity: a case study in Brazilian savannas

**DOI:** 10.1101/2025.06.23.661159

**Authors:** Cíntia Fernanda da Costa, Christhian Santana Cunha, Maria João Ramos Pereira

## Abstract

Wildfires have become a significant environmental challenge, exacerbated by climate change. Remote sensing technologies have advanced, but their use for monitoring fire frequency and severity remains limited, particularly in resource-constrained regions. To address this, we developed an open-source tool on Google Earth Engine that generates fire regime maps, combining fire frequency and severity data, thus allowing a temporal assessment of fire severity. The tool utilizes freely available satellite data, including Mapbiomas Fire data (1985-2020) and fire spectral indices (NBR, dNBR) from Sentinel-2A imagery (2017-2023). Our methodology provides high temporal and spatial resolution, enabling continuous analysis of fire patterns over time. It offers a cost-effective, scalable solution for monitoring fire dynamics, identifying areas most impacted by wildfires, and supporting informed decision-making. This capability helps prioritize conservation actions. The main advantages of this approach are its accessibility, adaptability, and processing time. This tool represents a significant advancement in remote sensing for conservation, aligning with global strategies to mitigate the impacts of climate change. It provides a practical, efficient, and replicable solution that not only supports local management efforts but also contributes to broader scientific and policy initiatives aimed at wildfire mitigation and sustainable ecosystem management.

## 1 INTRODUCTION

Fire is a natural event in many ecosystems worldwide, often playing a crucial ecological role. Savannas, grasslands, Mediterranean shrublands, tropical dry forests, heathlands and peatlands are some ecosystems where fire occurs naturally (Bond and Keeley 2005, Bowman et al. 2009, Pausas and Keeley 2009). Fire is a powerful eco-evolutionary force, influencing a wide range of processes; it shapes organismal traits, regulates population sizes, mediates species interactions, and determines community composition (McLauchlan et al. 2020). Fire also plays a key role in carbon and nutrient cycling, ultimately driving ecosystem function. However, even in fire-prone ecosystems, increasingly destructive wildfires driven by criminal activities or human-induced landscape changes demand urgent attention. Measuring fire severity is crucial for understanding its impact on ecosystems and guiding effective management strategies. Fire severity influences vegetation recovery, soil stability, and the redistribution of carbon and nutrients, which are essential for maintaining ecosystem function (Bowman et al. 2009, Salgado et al. 2024). High-severity fires can disrupt natural cycles, leading to biodiversity loss, soil degradation, and altered water dynamics, while low-severity fires often support ecological processes in fire-adapted systems (Kelly et al. 2023). Quantifying fire severity allows researchers and land managers to assess the ecological consequences of fires, evaluate post-fire recovery, and develop strategies to mitigate risks associated with both intense wildfires and fire suppression in fire-dependent ecosystems.

Fire is a natural ecological event typical of Brazilian savannas, occurring periodically and shaping the structure and dynamics of the local biota (Behling et al. 2007, Coutinho 1982, Arruda et al. 2016). However, land use changes, mostly resulting from the conversion of native vegetation caused by large-scale agriculture, mineral extraction and livestock production (Souza et al. 2020), provide opportunities for large fires to occur in these environments and also in typically forested areas. In addition, the relaxation of environmental inspection and financing policies in Brazil has facilitated irresponsible practices and rhetoric that weaken environmental protection, leading to an increase in criminal fires (Schmidt and Eloy 2020). In South America, Brazil stands out for having the highest incidence of wildfires (Li et al. 2020, White 2019), with at least 19.6% of its territory having experienced burning at least once from 1985 to 2020 (Alencar et al. 2022). During this period, most wildfires occurred in the Amazon and Cerrado ecoregions, while the Pantanal registered the highest proportion of the burned area. In most of these fires, native vegetation was significantly affected by the flames (Alencar et al. 2022). Unlike the Amazon, which is primarily forested, the Cerrado and Pantanal encompass a mosaic of landscapes, including predominantly savannas and open areas, as well as forested areas (Eiten 1972, Ribeiro and Walter 1998, Ivory et al. 2019). These ecosystems are largely classified as fire-dependent due to the array of adaptations that their plant and animal species have evolved (Berlinck and Batista 2020, Durigan 2020). Changes in fire intensity, spatial extent and periodicity can significantly impact biodiversity, directly and indirectly, at different spatial and temporal scales (Frizzo et al. 2011).

The Cerrado stands out for its exceptional biodiversity, surpassing all other savannas worldwide in the variety of plant and animal species (Mittermeier et al. 1999, Myers et al. 2000). However, by 2020, it is estimated that it has lost 50% of its native vegetation cover (Souza et al. 2020). The Pantanal, a large seasonal floodplain located in the heart of South America, covers 1.8% of Brazil’s territory (Souza et al. 2020), making it the largest floodplain in the world. Climatic changes associated with land use and cover alterations in the Cerrado include rising temperatures and reduced air humidity (Hofmann et al. 2021). Additionally, the decline in precipitation, primarily linked to shifts in atmospheric circulation caused by global climate change, affects both the Cerrado and the Pantanal. This decrease in rainfall is particularly pronounced during the dry season (june-september) and the onset of the rainy season (october-november), periods that coincide with the majority of wildfires (Correa et al. 2022, Shimabukuro et al. 2020, Alencar et al. 2022). While both ecoregions exhibit seasonal flooding patterns, the Cerrado is predominantly situated in elevated areas with well-drained soils that do not experience flooding (de Oliveira et al. 2023). In contrast, the Pantanal is subject to significant climate shifts, particularly reductions in rainfall and river flow (Araujo et al. 2018). Intense flooding periods foster the development of biomass-accumulating plants, and when the dry season arrives, this vegetation tends to become fuel, facilitating the spread of large fires in the region (Arruda et al. 2022, Marques et al. 2021). The incidence of wildfires increased significantly in 2020 (Pivello et al. 2021), marking the largest series of fires in the Pantanal in the last 20 years. During august and september, the Pantanal recorded approximately 22,000 fire outbreaks (INPE 2023). Nearly 30% of the total area of the Pantanal was consumed by fire, with an estimated minimum of 17 million vertebrates perishing as a direct consequence of those fires (Tomas et al. 2021). The continuous conversion of native areas, combined with climatic factors, is altering the seasonal dynamics of these ecosystems, prolonging dry periods and increasing the risk of wildfires in the future.

Affected areas show a distinct spectral response that can be monitored with remote sensing data (Pereira et al. 1999). In this context, several platforms are dedicated to monitoring the Brazilian territory through satellite imagery. The Fire Database (BDQueimadas- http://queimadas.dgi.inpe.br/queimadas) is the official monitoring system that provides data used by the Brazilian government to demonstrate its compliance with international agreements. It is an important tool for identifying and monitoring fire outbreaks, enabling the anticipation of risks in vegetative areas. The TerraBrasilis Portal also offers resources for querying, analyzing, and disseminating geographical data generated by monitoring projects such as PRODES and DETER (Assis 2019), all developed and maintained by the National Institute for Space Research (INPE). In addition, the MapBiomas Fire Project (Alencar et al. 2022) makes a significant contribution by validating these data and mapping fire scars across Brazilian territory from 1985 to 2022, with data captured at both monthly and annual intervals (Alencar et al. 2022). This comprehensive mapping effort is complemented by the Google Earth Engine, a cloud-computing platform developed by Google specifically designed to analyze and process of extensive geospatial datasets (Gorelick et al. 2017).

Recent studies have employed spectral indices, such as Normalized Burn Ratio (NBR) and Difference Normalized Burn Ratio (dNBR), to detect changes, pre and post-disturbance and, quantify wildfire occurrence, frequency, and severity (Key and Benson 2006). The NBR index is suitable for detecting changes in the landscape affected by fire and is considered effective in identifying burned areas (dos Santos et al. 2020). Variations in near-infrared (NIR) reflectance can indicate changes in photosynthetically active vegetation, while changes in shortwave infrared (SWIR) reflectance can signal reduced photosynthetically active vegetation, moisture content, ash deposition, and increased soil exposure. In studies related to forest fires in conservation units, it has been observed that the dNBR enhances the changes between NBR images, highlighting the presence of fire (dos Santos et al. 2020). The dNBR is obtained by subtracting the post-fire NBR image from the pre-fire NBR image (Roy et al. 2006).

Despite advancements in remote sensing, these tools have not been extensively used in the Brazilian context to analyze historical time series and discern the frequency and severity of fires. As a result, our primary aim was to develop an open-source code within Google Earth Engine to generate a temporal severity analysis map, capable of discerning areas with varying levels of severity and fire occurrence. We aimed to provide a robust, efficient, replicable, and accessible approach for monitoring fire frequency and severity, offering quantifiable, spatially explicit data with immediate applicability worldwide for environmental management and public policy. To accomplish this, we applied wildfire indices, NBR and dNBR, to map affected areas and understand the extent of the phenomenon in two Brazilian savanna regions. This analysis encompasses both fire severity, obtained through Sentinel-2A (2017-2023) satellite imagery, and fire frequency, using data from MapBiomas Fire Project (1985 – 2020, Alencar et al. 2022).

## 2 METHODS

### 2.1 FOCAL AREAS

Our focal areas were two conservation units in Brazil: Chapada dos Guimarães Environmental Protection Area (APA), in the Cerrado ecoregion and the SESC Pantanal Private Natural Heritage Reserve (RPPN), situated in the Pantanal ecoregion (Figure 1). Both are located in the state of Mato Grosso and have a history of fires, with fire management strategies outlined in their respective management plans.

**Figure 1.**
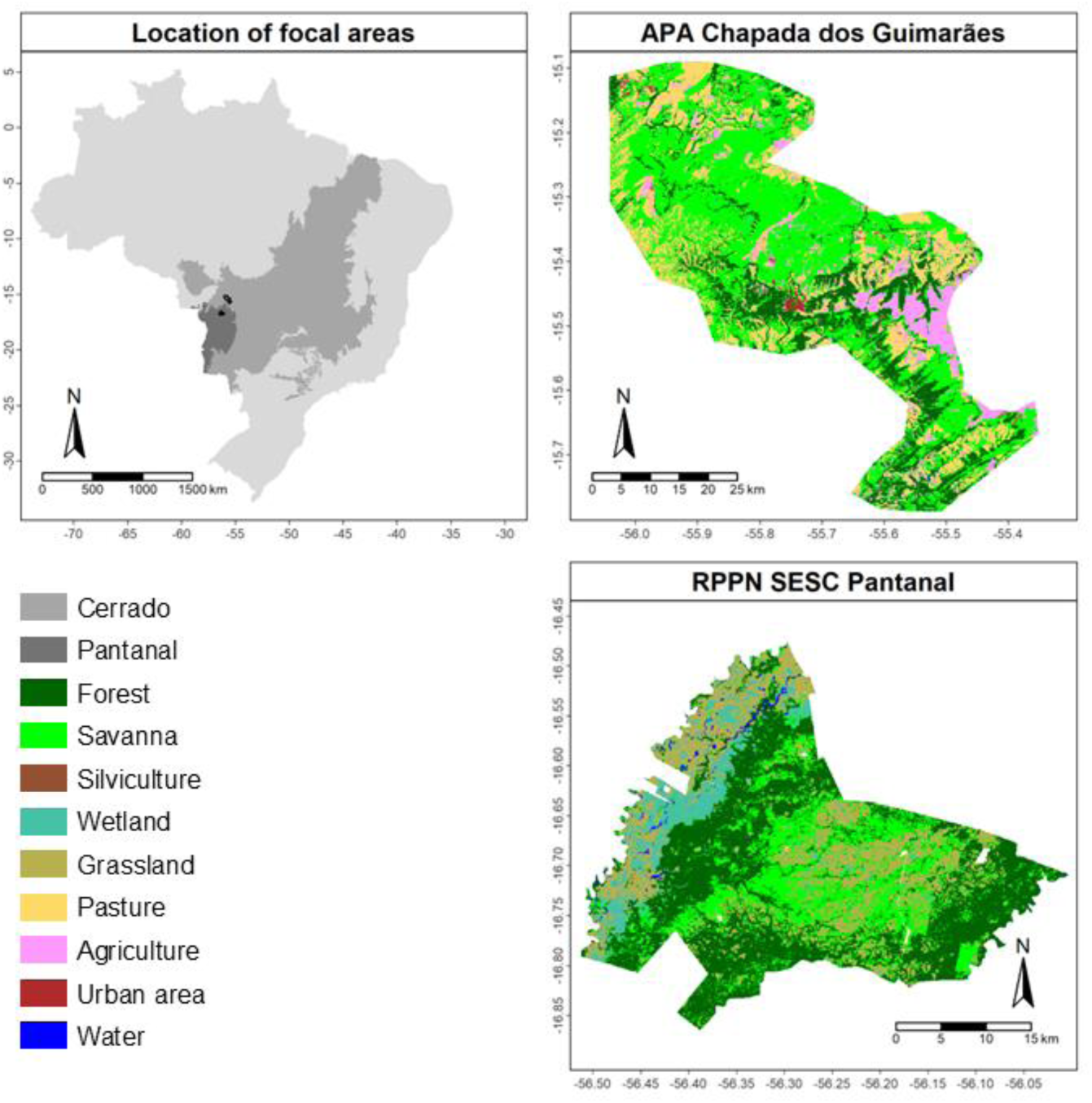
Map showing Brazil in light gray, with the Cerrado ecoregion in medium gray and the Pantanal ecoregion in dark gray, indicating the two sampling areas. Also included are detailed maps of land use and land cover for the Chapada dos Guimarães Environmental Protection Area (APA) and the SESC Pantanal Private Natural Heritage Reserve (RPPN), for 2023 (MapBiomas Project 2022).

#### 2.1.1 CHAPADA DOS GUIMARÃES ENVIRONMENTAL PROTECTION AREA (APA)

The protected area extends over 251,848 hectares and consists of a set of conservation units, such as the Chapada dos Guimarães National Park (PARNA), the Cuiabá-Chapada dos Guimarães-Mirante Park Road, the Quineira State Park, and the Natural Monument of the Geodesic Center of Latin America. The area features well-preserved ecosystems, including riparian forests, gallery forests, dry forests, and Cerrado *stricto sensu*, as well as savanna formations, palm swamps, and grassland systems (Eiten 1972, Ribeiro and Walter 1998). Additionally, the protected area features sections that have been converted into pastures and agricultural lands, inherited from before the implementation of its designation in 2002 (Law No. 7,804/02) (Figure 1). The area is influenced by the rainy season, from october to march, is followed by dry periods from april to september, resulting in an average annual rainfall of 1500 mm (ICMBio 2009). One of the region’s most significant features is its preservation of vital water resources, including springs, rivers, and recharge areas of the Guarani Aquifer, one of the world’s primary freshwater reserves (Aráujo et al. 1995).

#### 2.1.2 SESC PANTANAL PRIVATE NATURAL HERITAGE RESERVE (RPPN)

Considered the largest Private Natural Heritage Reserve in Brazil, spanning 108,000 hectares in the municipality of Barão do Melgaço, Mato Grosso, the RPPN is located amidst extensive areas of relatively preserved native vegetation, with some pastoral areas (Figure 1). The region is characterized by a tropical climate with dry winters and rainy summers, receiving annual rainfall ranging between 1000 mm and 1600 mm (Hofmann et al. 2010). The high biodiversity of the Pantanal results from the integration of flora and fauna elements from the Cerrado, Chaco, Amazon, and Atlantic Forest. Additionally, the diversity of habitats in the Pantanal is closely linked to geomorphological variations and the unpredictability of flooding, which creates distinct environments: permanently flooded areas (such as forests dominated by Cambará trees), rarely flooded areas (such as fields with murundus - elevated mounds in floodplain landscapes), non-flooded areas (such as forests with acuri palm trees), and zones with annually variable dynamics (Ivory et al. 2019, Russell et al. 2003, Cunha et al. 2007). The flooding cycle also annually replenishes soil nutrients, boosting ecosystem productivity and supporting unparalleled population abundance (Catian et al. 2019).

### 2.2 OPEN-SOURCE CODE AND OUTPUTS

The code supporting this study is officially hosted on GitHub, ensuring transparency, reproducibility, and accessibility for future validation and collaboration. Interested researchers and professionals can access it through the following link: https://github.com/Kejorureu/Temporal-assessment-of-fire-severity. The stages of code development and the final outputs are discussed in detail in the following sections. The methodological process is depicted in the flowchart in Figure 2.

**Figure 2.**
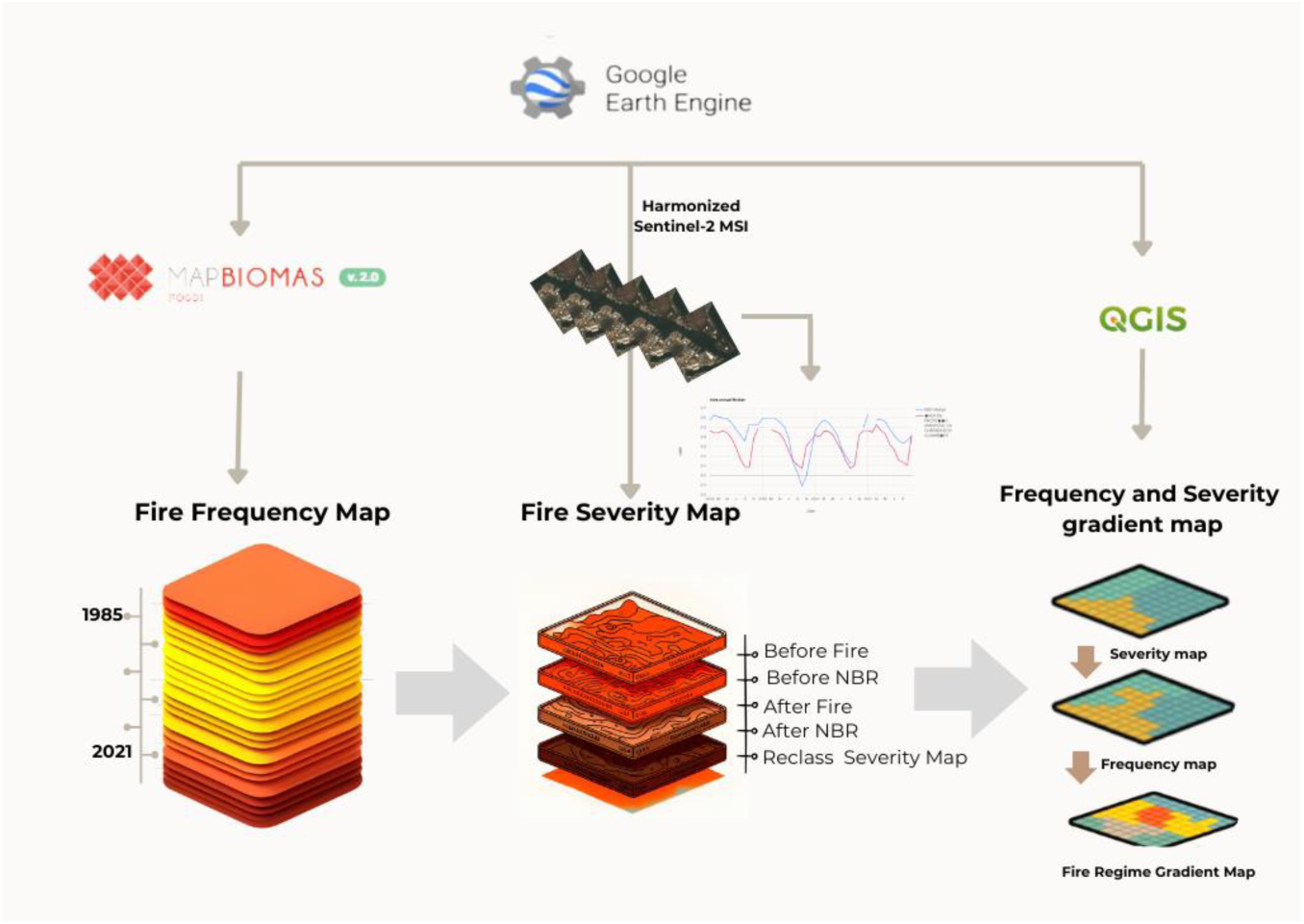
Flowchart of the activities and processes carried out for the development of the fire regime gradient map, representing the temporal assessment of fire severity for the Chapada dos Guimarães Environmental Protection Area (APA) and the SESC Pantanal Private Natural Heritage Reserve (RPPN), Mato Grosso, Brazil.

#### 2.2.1 FIRE FREQUENCY MAP

Using data sourced from the Mapbiomas Fire Project (Alencar et al. 2022), accessed through the Google Earth Engine via toolkit, we created a map depicting the frequency of wildfire events. The term ‘fire frequency’ denotes the number of occurrences a specific location experiences fires within a defined timeframe. We processed a collection of images containing annual burned area and fire frequency data. For the fire frequency analysis from 1985 to 2020, we assigned a binary value to each pixel, 1 if it burned during the year and 0 if it did not. We then summed these values across years, normalized the result for each pixel to a scale of 0 to 1, and multiplied by 100 to express the values as percentages, converting the final results to integers. We then summed all the annual layers into a single layer. The *fire frequency* product enabled us to estimate the recurrence of events in the studied pixels. Subsequently, we classified the fire frequency data into six distinct categories: No fire record, low fire record, moderate-low fire record, moderate fire record, moderate-high fire record and high fire record (Figure 3).

**Figure 3.**
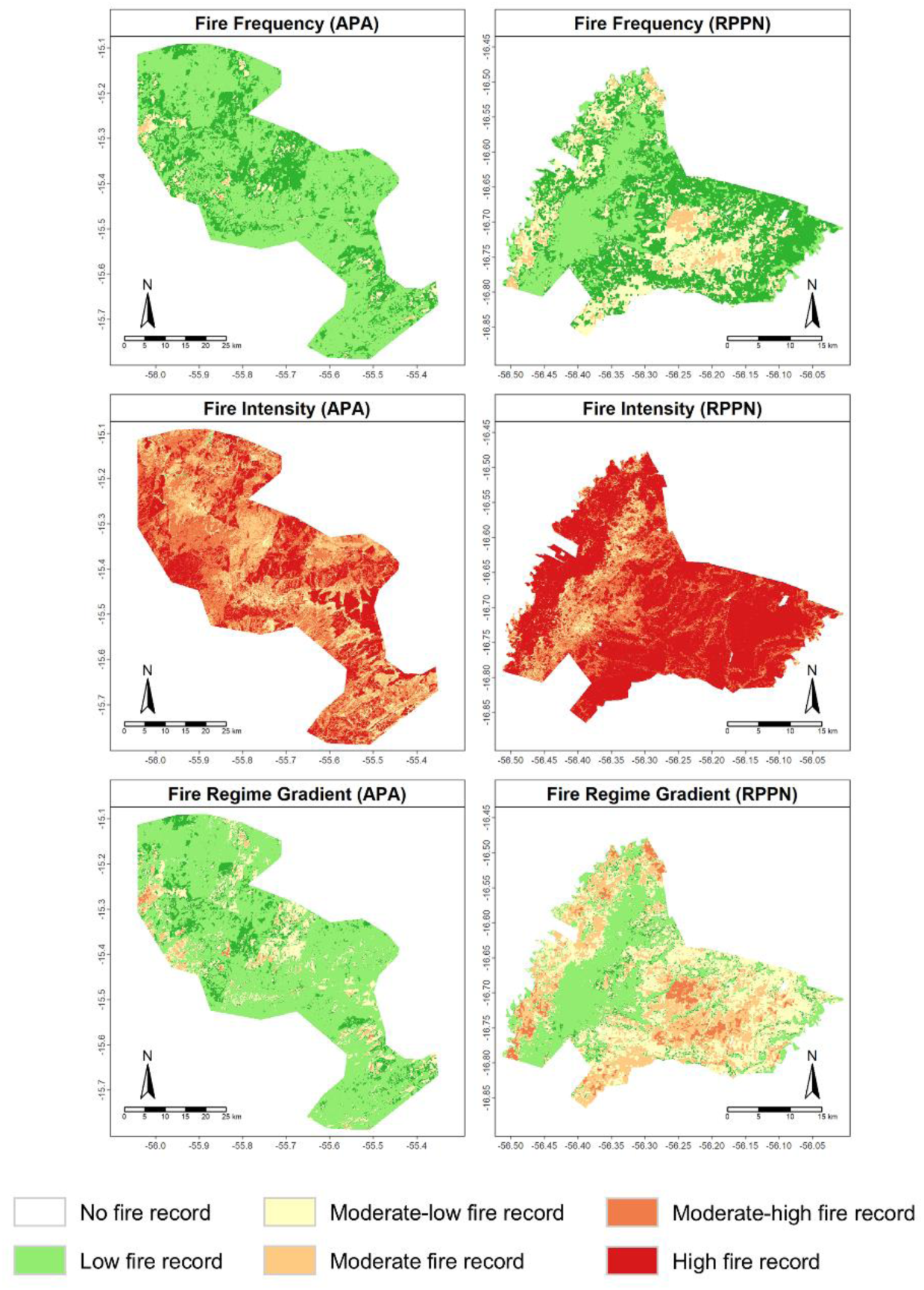
Final products developed for the Chapada dos Guimarães Environmental Protection Area (APA) and the SESC Pantanal Private Natural Heritage Reserve (RPPN) in Brazil. For fire frequency, we used Collection 2 of the annual MapBiomas Fire Project (1985–2022, Alencar et al. 2022). Fire intensity maps were generated using Sentinel-2A imagery (2019–2022) and reclassified based on the Differenced Normalized Burn Ratio (dNBR) method, as proposed by the United States Geological Survey (USGS). The intensity classes are as follows: values between -0.1 and 0.1 indicate unburned areas; values between 0.1 and 0.27 represent Low intensity; values between 0.27 and 0.44 correspond to Moderate-low intensity; values between 0.44 and 0.66 indicate Moderate-high intensity; and values greater than 0.66 represent high intensity. Additionally, fire regime gradient maps combining frequency and intensity data were produced.

#### 2.2.2 FIRE SEVERITY MAP

Fire severity quantifies the energy released during a fire, typically measured as heat per unit area. Our analysis of wildfire indices utilized Sentinel-2A imagery from the ‘COPERNICUS/S2_SR_HARMONIZED’ collection, covering the period from March 2017 to January 2023. This high-resolution multispectral imaging mission, conducted by Copernicus Sentinel-2A, provides extensive coverage to support land monitoring studies. The mission enables the observation of various environmental factors, including vegetation, soil, water cover, inland water bodies, and coastal regions, with a revisiting time of five days. To identify burned areas, we applied the Normalized Burn Ratio (NBR) index, following the methodology described by Key and Benson (2006).

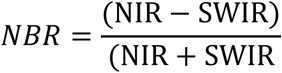

High NBR values indicate healthy, dense vegetation, while values near zero reflect exposed soil or sparse vegetation with lower NIR and SWIR reflectance. Negative values indicate the presence of water, which reflects less in the NIR. In the context of fires, areas with reduced NBR values post-fire indicate degraded vegetation and exposed surfaces. The next step of the code then involves applying the addNBR function within our designated areas of interest, leveraging the NIR (B8) and SWIR-2 (B12) bands from Sentinel-2A imagery. Furthermore, we used the maskCloudAndShadowsSR function to mask out clouds and shadows present in the images. Following the definition of the image collection for computing and interpreting the indices, we advanced to the image processing phase. This entailed filtering based on date, location, and the proportion of cloudy pixels. We executed the cloud and shadow masking function, chose pertinent bands, and computed the NBR. Subsequently, we merged the images by date using the mosaicByDate function. We reduced the data to a monthly resolution and generated a time series graph of the intra-annual median NBR (Figure 4).

**Figure 4.**
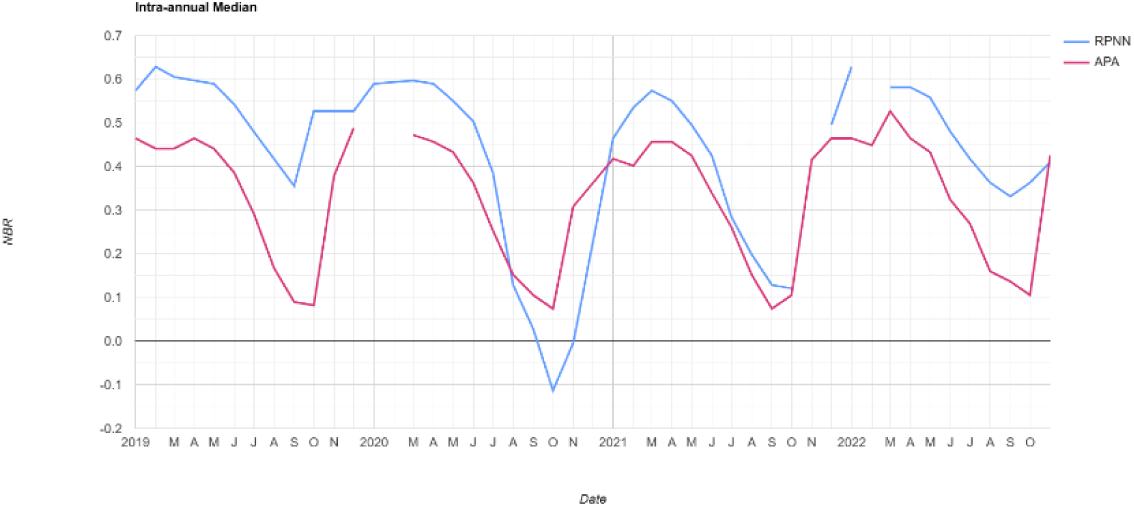
Graphical representation of the intra-annual median time series of the Normalized Burn Ratio (NBR) for the Chapada dos Guimarães Environmental Protection Area (APA), depicted in red, and the SESC Pantanal Private Natural Heritage Reserve (RPPN), in blue, both located in Mato Grosso, Brazil.

When we analyzed the time series graph of the NBR for the conservation units, we noticed the presence of seasonalities with both positive and negative NBR values (see Figure 4). We decided to select the period with the lowest NBR (negative NBR) to define the post-fire area and an image with a higher NBR for the pre-fire period (positive NBR). The calculation of pre-NBR and post-NBR enables the study of change detection through multi-temporal visual analysis. In other words, it becomes possible to compare the alterations occurring between a period before and after the event (Supplementary Figure 1). Subsequently, we used the dNBR, to identify changes between pre-fire and post-fire periods.

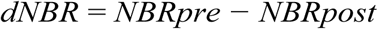

After calculating the dNBR, we reclassified the pixels using the methodology outlined by the United States Geological Survey (USGS). The following classes were adopted: Unburned, Low intensity, Moderate-low intensity, Moderate-high intensity, and High intensity (Figure 3).

#### 2.2.3. FIRE REGIME GRADIENT MAP

After analyzing the previous mappings, the intensity and frequency of wildfire data were harmonized in the QGIS program (QGIS Development Team 2022), following the provided classification: No fire record, low fire record, moderate-low fire record, moderate fire record, moderate-high fire record and high fire record (Figure 3). This process resulted in a fire regime gradient map, which allows identifying areas with frequent and intensity fire occurrences, indicating, for example, more vulnerable and impacted locations. It serves as a tool for assessing historical fire patterns and guiding management in the protection, management, and restoration of these areas.

## 3 RESULTS AND DISCUSSION

### 3.1 OPEN-SOURCE TOOL FOR ANALYZING TEMPORAL FIRE SEVERITY USING GOOGLE EARTH ENGINE

Our open-source code, available on Google Earth Engine, provides a robust tool for analyzing temporal patterns of fire severity in any region of interest. Due to its high adaptability, it only requires the substitution of the area of interest to initiate the analysis. Users can achieve personalized results for their study areas by making minimal adjustments to the cloud cover percentage, which varies by region. The flexible adaptation of the code also allows it to be integrated with other geospatial databases, such as land use change maps, enhancing its applicability in multidimensional studies. For example, it is possible to correlate fire patterns with soil and vegetation health indicators, providing a broader understanding of the ecological impacts of wildfires. This multidisciplinary approach is particularly relevant in the context of climate change, where data-driven adaptive management is essential. Savannas and other regions worldwide are experiencing rapid changes in land use (Williams et al. 2022). Understanding how these transformations affect, or are affected by, the occurrence and intensity of fires is crucial for interpreting how species and ecosystems respond to these pressures. Through time series analysis of fire patterns and landscape changes, we identified areas with a higher likelihood of fire occurrence and greater vulnerability to loss. This enables a more efficient allocation of resources for fire prevention and response efforts, focusing on strategic areas for monitoring, prevention, mitigation, and future actions.

Our product has been developed to assist both researchers and decision-makers. It can serve as a foundation for identifying the most vulnerable sites that require greater attention in planning-controlled fire management and developing fire prevention actions. From the identified patterns and critical areas, it is possible to establish sampling grids ranging from low to high intensity and frequency of fire regime. This allows for continuous monitoring of fire behavior in the areas of interest to understand how the effects of fire impact the populations and communities of flora and fauna in these locations.

Sentinel-2A stands out for its high spatial resolution, with visible, near-infrared (NIR), and shortwave infrared (SWIR) bands available at 10, 20, and 60 meters. This makes it ideal for detecting detailed patterns in land use and cover, such as assessing wildfire severity. Additionally, its revisit interval of five days, enabled by the operation of two satellites, allows for near-continuous monitoring and increases the likelihood of acquiring cloud-free images. Sentinel-2A also offers a wide range of spectral bands, including the Red Edge band, which is particularly useful for advanced vegetation and soil analyses. For these reasons, and considering the focus of our case study, we selected Sentinel-2A. However, users of the code can choose longer time series or access specific information from previous years using Landsat satellite data. The choice between Sentinel-2A and Landsat satellites for remote sensing analyses should consider the project scope, as well as the required temporal and spatial scales, by evaluating the strengths and limitations of each sensor.

Landsat provides a long record of Earth observation, with data available since 1972, enabling historical studies and long-term analyses of environmental changes. While its spatial resolution of 30 meters is lower, it remains suitable for regional-scale studies. However, its revisit frequency of 16 days can limit image availability in areas with frequent cloud cover. To use Landsat, the data processing code must be adjusted. This involves switching the data source to specific collections, such as Landsat Collection 2 Tier 1 or Tier 2, and adapting spectral indices to the corresponding bands, as their nomenclature differs from Sentinel-2A. For example, Sentinel-2A’s band 8 (NIR) corresponds to band 5 in Landsat 8. Additionally, steps in the code that depend on specific resolutions or revisit frequencies must be modified to ensure compatibility. The lower revisit frequency of Landsat may require the use of composite data over longer periods to compensate for reduced image availability. Thus, the decision between Sentinel-2A and Landsat depends on the study’s priorities, and it is essential to adapt analytical tools to maximize the advantages of each sensor.

### 3.2 TEMPORAL ASSESSMENT OF FIRE SEVERITY IN BRAZILIAN SAVANNAS

Our assessment of the fire frequency maps revealed overlaps during the analyzed period (1985-2020), allowing us to identify the frequency at which the areas were affected by wildfires. The frequency of those fires varied, ranging from 1 to 5 times for the Chapada dos Guimarães Environmental Protection Area (APA) and from 1 to 4 times for the SESC Pantanal Private Natural Heritage Reserve (RPPN). In the case of the Chapada dos Guimarães Environmental Protection Area, our analysis showed that ca. 68% of the total burned areas experienced fire once, while approximately 30% of the area suffered two or three fire occurrences. Only a small fraction, less than 1%, was affected by fires 4 to 5 times during the analyzed time frame. In this area, there was a higher recurrence of fires in pastureland followed by savanna formations. As for the SESC Pantanal Private Natural Heritage Reserve (RPPN), the results indicated that most burned areas experienced a fire at least twice, making up 42% of the total. Around 26% of the areas were burned once or three times, while 5% were affected by fires on four occasions. In this area, fires impacted native grasslands and savanna formations with greater severity. It is also noteworthy that significant wetland areas were burned three times, while forested areas experienced fires twice, although in much smaller proportions.

In terms of intensity, the fires varied from low to high intensity, impacting each pixel at least once in both areas. Our findings reveal that moderate to high-intensity fires affected 70% of the Chapada dos Guimarães Environmental Protection Area (APA) and 90% of the SESC Pantanal Private Natural Heritage Reserve (RPPN), spanning all land use and land cover classes within these protected areas. These results highlight a significant effect on both areas, with over 155,000 hectares suffering high-intensity fires. It is crucial to highlight that this timeframe encompassed the wildfires of 2019 and 2020, which were the largest in decades in both areas. These events were truly unprecedented and stood out from the rest of the fire history (Pivello et al. 2021).

The data from the fire regime gradient map indicated that only 1% of the Chapada dos Guimarães Environmental Protection Area (APA) and 5% of the SESC Pantanal Private Natural Heritage Reserve (RPPN) were impacted by high-intensity and high-frequency fires. Conversely, approximately 68% of the Chapada dos Guimarães Environmental Protection Area (APA) was affected by low-frequency and low-intensity fires. The severity regime of fire in this area suggests a regular occurrence of such events in this region. Generally, ecosystems adapted to fire develop the ability to recover from regular fire regimes (He et al. 2019). However, it is noteworthy that a small part of the area exhibited moderate or moderate-high intensity regimes, particularly in pastureland.

This is particularly concerning, as recent estimates from MapBiomas show that between 1985 and 2022, the conversion of native vegetation in the Cerrado for agriculture and pastureland increased by 6.2 times (Knapp et al. 2009). These land use and land cover changes, coupled with more frequent and intense droughts and reduced rainfall (Souza et al. 2020, Hofmann et al. 2023), highlight the urgent need for careful fire management in the region to prevent devastating wildfires, such as those experienced in 2019 and 2020.In the specific context of the Chapada dos Guimarães National Park (PARNA), located within the protected area, where most of the native vegetation is preserved, prescribed burns are conducted annually. These activities are typically supervised by firefighting agencies, municipal authorities responsible for public management, professionals, and park managers. The primary objective of prescribed burns is to reduce the accumulation of combustible plant material, control the growth of invasive plants, promote the regeneration of native vegetation, and mitigate the risk of uncontrollable fires (Francos and Úbeda 2021, Knapp et al. 2009). Consequently, this approach has the potential to significantly contribute to the preservation of biological diversity and ecological processes in the protected area, maintaining the fire regime within the regular conservation unit. However, maintaining ecosystem balance, reducing the risk of uncontrollable wildfires, and promoting the regeneration of native vegetation while minimizing long-term ecological damage relies on carefully planned prescribed fires—an approach that is not always effectively implemented. Like Chapada dos Guimarães National Park (PARNA), the RPPN SESC Pantanal (RPPN) also maintains a longstanding fire monitoring and combat program. Our data revealed that this area experienced approximately 62% impact from fires of moderate intensity and frequency, with some areas showing moderate to high-intensity and frequency. Moderate-intensity fires represent a complex combination of surface and crown fires, where flames spread both through the lower vegetation layer and the tree canopy (Halofsky et al. 2011). On the other hand, high-intensity fires are characterized by vigorous and rapid combustion, consuming large amounts of vegetation within a short period (Rego et al. 2021). These fires tend to spread extremely quickly, fueled by favorable weather conditions such as terrain features, strong winds, and low humidity (Huang and Gao 2021). As a result, they can cause substantial damage to the landscape, resulting in significant loss of native vegetation and animal communities, habitats, and other natural resources, as well as posing a serious risk to human communities living in vulnerable areas (Nawaz and Henze 2020, Sant’Anna and Rocha 2020). Consequently, the recovery, if possible, of those areas may require a substantially longer time. Moreover, the vegetation may not have enough time to fully regenerate between fires, which can lead to changes in the structuring of plant and animal communities over time (Ross et al. 2023, Forney and Peacock 2024).

Over the past decades, the Pantanal has been severely affected by land use conversion, with a significant expansion of areas allocated for pasture (Souza et al. 2020). Between 1985 and 2022, pastureland expanded threefold, leading to a net loss of 10% of native vegetation in the region (Souza et al. 2020). This pattern of environmental degradation is already apparent in the areas surrounding the SESC Pantanal Private Natural Heritage Reserve (RPPN). A stark example occurred in April 2024, when a single farmer invested five million dollars to clear 80,000 hectares of native forest for cattle pasture in northern Pantanal, near the reserve. This widespread deforestation has been accompanied by unsustainable agricultural practices (Marques et al. 2021, Guerra et al. 2020), including the indiscriminate use of pesticides (Viana et al. 2023, de Oliveira and Roque et al. 2021). In the abovementioned case, the farmer applied 25 different pesticides, including 2,4-D, a component of the notorious Agent Orange herbicide. These practices not only threaten local biodiversity but also pose significant risks to the health and safety of local human communities and those who depend on the Pantanal’s natural resources for their livelihoods (Bonner and Alavanja 2017).

## 4 CONCLUSIONS

Our tool offers a robust, efficient, and replicable approach for assessing fire severity through time in any area of interest, with significant implications for both environmental management and public policy. Key expected outcomes include high temporal and spatial resolution, enabled by Sentinel-2A imagery. This allows for detailed, continuous analysis over a five-year period, helping to identify critical temporal and seasonal patterns in fire occurrences. Such insights are essential for understanding fire dynamics in sensitive ecosystems and informing effective conservation strategies. The innovative feature of this methodology is its automation and accessibility. By leveraging open-source tools and freely available satellite imagery, it democratizes environmental monitoring, offering a cost-effective solution for protected area managers, even in resource-limited settings. Furthermore, its open-source nature encourages collaboration among different user groups and the continuous improvement of the tool. This broadens its applicability, making it viable for use by various stakeholders at both regional and global scales. Additionally, the methodology excels in identifying priority conservation areas, and generating dNBR and severity maps that provide valuable data for management actions, such as habitat restoration or fire prevention. This capability supports optimized resource allocation in conservation efforts.

Moreover, the data generated with our tool can inform public policies aimed at mitigating the environmental impacts of fires. By providing clear, quantifiable, and spatially explicit information, it supports informed decision-making and helps develop sustainable management strategies for vulnerable ecosystems. The rapid detection of fire events and near-real-time severity classification further enhances the ability to implement timely interventions, reducing long-term environmental damage and fostering effective responses to fire threats. This tool aligns with global environmental monitoring agendas and contributes to more effective management in times of increasing climate challenges.

## Supporting information

https://github.com/Kejorureu/Temporal-assessment-of-fire-severity

## 7 FUNDING SOURCE ACKNOWLEDGMENTS

We would like to express our gratitude to the Rufford Foundation for supporting our project, Effects of Fire on Bat Diversity and Occupancy in the Megadiverse Brazilian Savannas, with a Small Grant (41558-2 ID). We also thank the Brazilian Fund for Biodiversity – FUNBIO (014/2023), Bat Conservation International (BCI) for the 2023 Student Research Scholarship, and the Brazilian Bat Research Society – SBEQ for the Small Grants in Biology, Ecology, and Bat Conservation. Our thanks to the Brazilian National Council for Scientific and Technological Development – CNPq for providing a productivity grant to MJRP. This work was supported by the Coordination for the Improvement of Higher Education Personnel – Brazil (CAPES) with a PhD scholarship for CC and CSC (Funding Code 001).

## 8 AUTHOR CONTRIBUTIONS

The authors contributed equally to the article and approved the submitted version.

## Notes

https://github.com/Kejorureu/Temporal-assessment-of-fire-severity

### Competing Interest Statement

The authors have declared no competing interest.

