## Supplementary material for "Temporal assessment of fire severity: a case study in Brazilian savannas": https://github.com/Kejorureu/Temporal-assessment-of-fire-severity

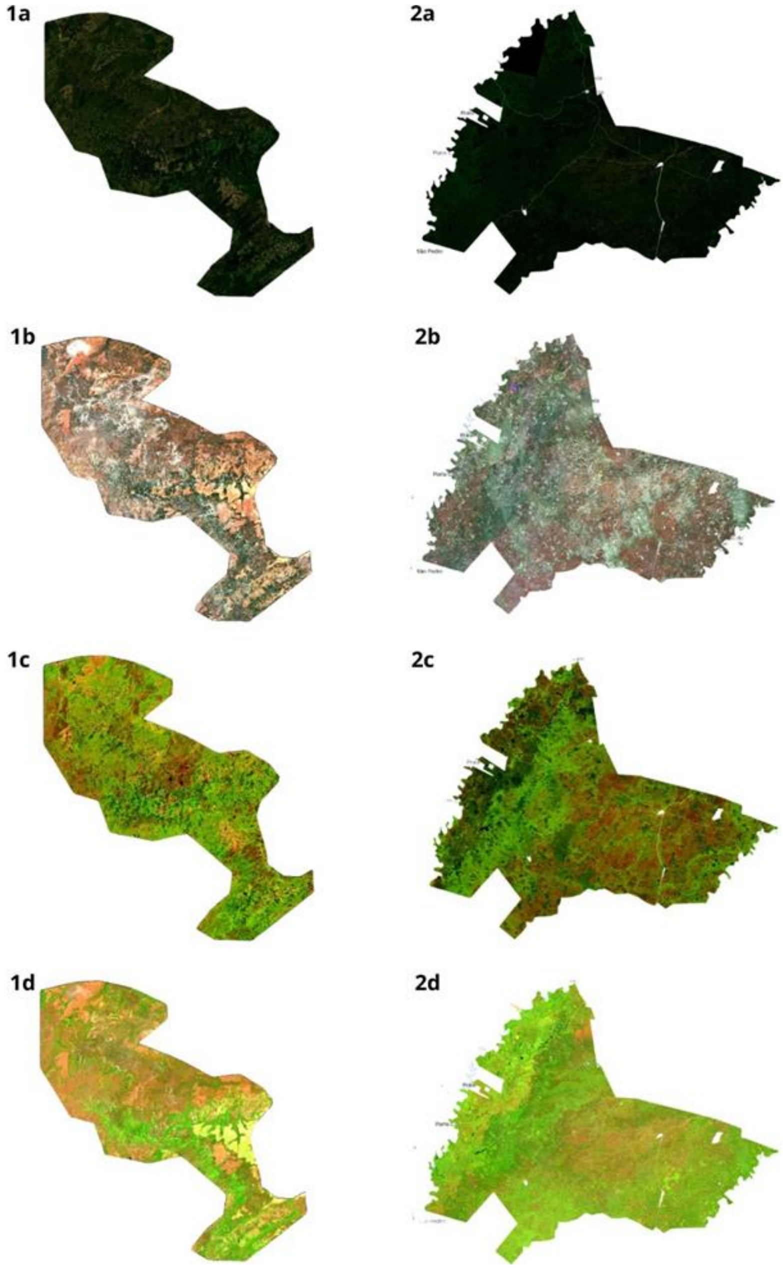


FIGURE 1: Sentinel-2 image mosaic representing: 1 for the Chapada dos Guimarães Environmental Protection Area (APA) and 2 for the SESC Pantanal Private Natural Heritage Reserve (RPPN). In a) Color composition (RGB) of the period with a low number or absence of fire events; b) Recording of fire-affected pixels during the analyzed period; c) and d) False-color compositions, composed of SWIR, NIR, and red bands.
